# insideOutside: an accessible algorithm for classifying interior and exterior points, with applications in embryology

**DOI:** 10.1101/2021.11.15.468285

**Authors:** Stanley E. Strawbridge, Agata Kurowski, Elena Corujo-Simon, Alastair N. Fletcher, Jennifer Nichols, Alexander G. Fletcher

## Abstract

A crucial aspect of embryology is relating the position of individual cells to the broader geometry of the embryo. A classic example of this is the first cell-fate decision of the mouse embryo, where interior cells become inner cell mass and exterior cells become trophectoderm. Fluorescent labelling, imaging, and quantification of tissue-specific proteins have advanced our understanding of this dynamic process. However instances arise where these markers are either not available, or not reliable, and we are left only with the cells’ spatial locations. Therefore, a simple, robust method for classifying interior and exterior cells of an embryo using spatial information is required. Here, we describe a simple mathematical framework and an unsupervised machine learning approach, termed `insideOutside`, for classifying interior and exterior points of a three-dimensional point-cloud, a common output from imaged cells within the early mouse embryo. We benchmark our method against other published methods to demonstrate that it yields greater accuracy in classification of nuclei from the pre-implantation mouse embryos and greater accuracy when challenged with local surface concavities. We have made MATLAB and Python implementations of the method freely available. This method should prove useful for embryology, with broader applications to similar data arising in the life sciences.

## Introduction

The mouse embryo undergoes three major morphogenetic events between fertilization and implantation: compaction, cavitation, and hatching (Fig. 1A,B) (Tarkowski and Wróblewska, 1967; Smith and McLaren, 1977; Yoshinaga et al., 1976). Compaction coincides with the first binary cell-fate decision, which is ultimately driven by cellular position within the embryo (Fig. 1C) (Tarkowski and Wróblewska, 1967). Exterior cells polarize to become the extraembryonic trophectoderm (TE), precursors of the placenta (Lawson et al., 1999), while interior cells become the inner cell mass (ICM) (Ziomek and Johnson, 1980; Johnson and Ziomek, 1981). ICM cells then undergo a second binary cell-fate decision to become either the embryonic epiblast, source of the foetus (Gardner and Rossant, 1979) and embryonic stem cells (Evans and Kaufman, 1981; Martin, 1981), or the primitive endoderm (PrE), founder of the yolk sac (Gardner and Johnson, 1972). This second cell-fate decision coincides with cavitation, where a fluid-filled cavity, called the blastocoel, forms between the TE and one side of the ICM (Smith and McLaren, 1977). Finally, prior to implantation, the embryo must hatch from the zona pellucida (Malter and Cohen, 1989).

**Figure 1:**
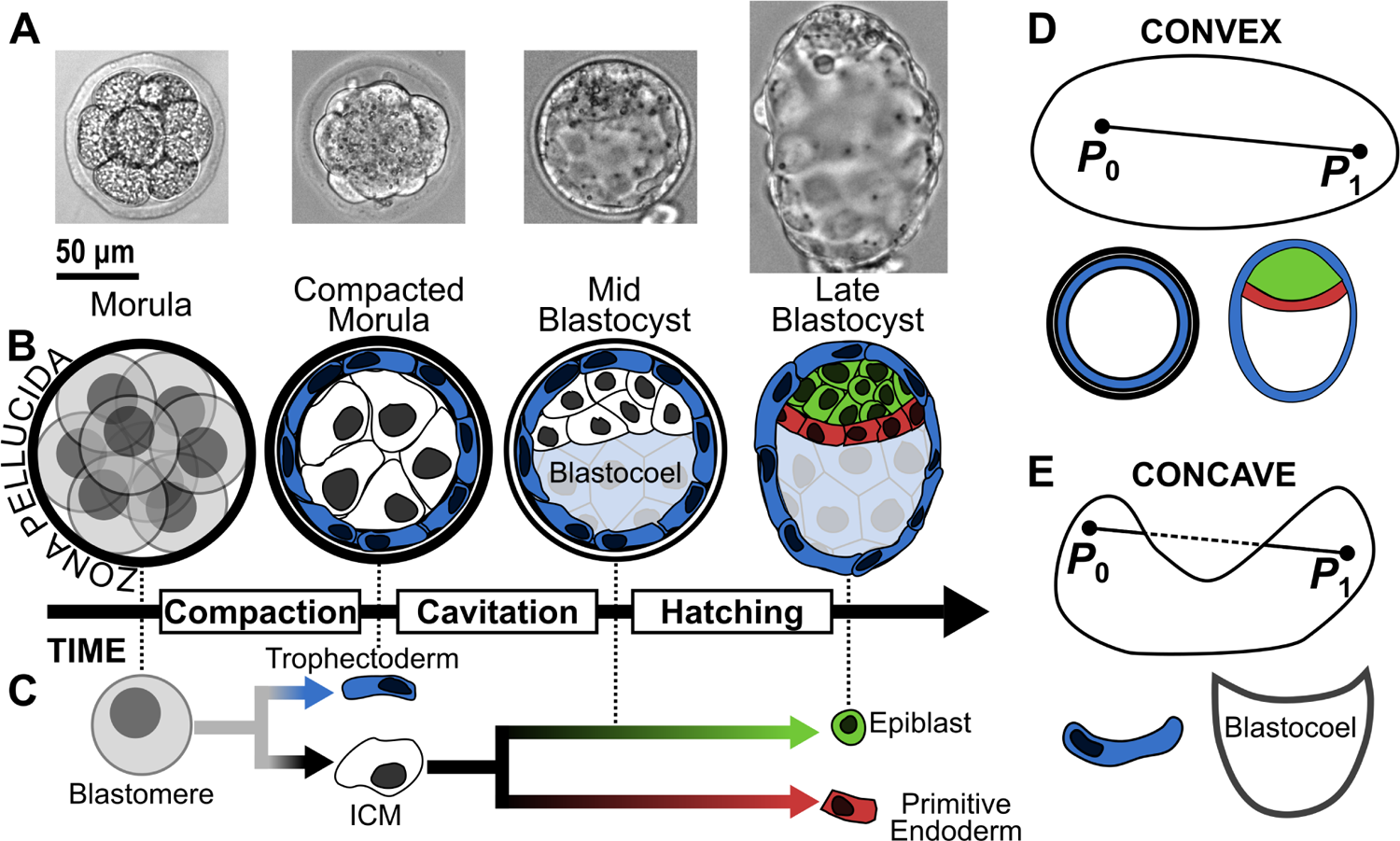
**A.** Bright-field images from pre-implantation mouse development, from morula to late blastocyst. **B.** Three major morphogenetic events occur during pre-implantation mouse development: compaction, cavitation, and hatching. **C.** Cells in the pre-implantation embryo make two sets of binary cell-fate decisions: first, blastomeres become inner cell mass (ICM) (interior) or trophectoderm (exterior); second, ICM become epiblast or primitive endoderm. These decisions coincide with compaction and cavitation, respectively, and are completed by hatching. **D.** A shape is convex if for any pair of points, *P*_0_ and *P*_1_, the resulting line segment is entirely contained within the shape; biological examples of convex shapes include the compacted morula and blastocyst. **E.** A shape is concave if there exists at least one pair of points whose resulting line segment passes to the exterior of the shape; biological examples of concave shapes include trophectoderm cells and the blastocoel cavity.

Molecular profiling of these tissues through RNA sequencing (Guo et al., 2010, 2017) and immunohistochemistry (Chazaud et al., 2006; Niwa et al., 2005; Palmieri et al., 1994) has revealed key lineage markers such as NANOG and GATA6. These lineage markers have been used to study the dynamic emergence and plasticity of distinct cell identities during pre-implantation development by employing fluorescent reporter knock-ins (Arnold et al., 2011; Grabarek et al., 2012; Hamilton et al., 2003; Kalkan et al., 2017; McDole and Zheng, 2012). However, in the mouse there remain instances where reliable lineage markers do not exist (Plusa et al., 2008) or cease to faithfully mark their lineage (Le Bin et al., 2014; Schrode et al., 2014; Bessonnard et al., 2014), while in other mammals such as humans and non-human primates, such lineage markers are not yet established (Boroviak et al., 2018; Guo et al., 2021; Stirparo et al., 2018). In such cases, we must find alternative methods to classify the tissues under investigation.

For decades, spatial information has been used to help classify cell populations in the pre-implantation mouse embryo (Fleming, 1987; Nichols and Gardner, 1984). Recent advances in image acquisition and processing technologies have improved the accuracy of this spatial information. A common analysis method is quantitative immunofluorescence (qIF) of cell nuclei, whereby three-dimensional (3D) confocal fluorescence microscopy images of nuclei are segmented and quantified using software such as Fiji (Schindelin et al., 2012), MINS (Lou et al., 2014), or Nessys (Blin et al., 2019). Output parameters from qIF include total nuclear fluorescence, nuclear volume, and the geometric centre (centroid) of the nucleus. The centroid point-cloud can then be used to classify individual nuclei by their relative positions. Classification of interior and exterior nuclei is of particular interest when investigating the relationship between the cells of the ICM (interior) and the TE (exterior).

To date, three methods have been used to classify interior and exterior nuclei of the mouse embryo from qIF. We refer to these as the random sample consensus (RANSAC) Ellipsoidal, Convex Hull, and `insideOutside` methods. The RANSAC Ellipsoidal method, employed by MINS (Lou et al., 2014), robustly fits an ellipsoid to the point-cloud generated by segmented nuclear centroids through the RANSAC iterative method (Fischler and Bolles, 1981). Each nucleus is then classified as exterior if the distance from the ellipsoid’s centre to the nuclear centroid exceeds 0.95 times the distance from the ellipsoid’s centre to the point on the ellipsoid that is closest to the nucleus’s centroid; otherwise, the cell is classified as interior. MINS has been widely used for qIF and has been cited in nearly 100 manuscripts. The Convex Hull method, employed by the spatial analysis software IVEN (Forsyth et al., 2021), constructs a convex hull, the smallest convex set that contains all centroids, from all nuclear centroids of the embryo and then classifies a nucleus as exterior if it belongs to the boundary of the convex hull. IVEN allows for manual correction of the classification, but this requires user input which may introduce bias. The `insideOutside` method, a preliminary version of which was employed by Stirparo et al. (2021), is an accessible position-based approach to the classification of interior and exterior nuclei.

A fourth method, which we refer to as the Naïve Ellipsoidal method, was introduced by Forsyth et al. (2021) for their benchmarking. This approach accepts the first fit of an ellipsoid to the point-cloud, as opposed to generating a consensus ellipsoid through iterative random sampling, as in the RANSAC approach. The same distance rule as the RANSAC Ellipsoidal method is then applied to classify cell positions.

These four methods share the common assumption that the embryo is convex. A shape is said to be convex if the line segment between any pair of points within the shape is entirely contained within the shape (Fig. 1D); otherwise the shape is said to be concave (Fig. 1E). The RANSAC Ellipsoidal method models the embryo as an ellipsoid, which is a convex shape. Similarly, the Convex Hull method explicitly defines the exterior points as being a member of a convex shape. Therefore, these methods underperform if the surface of the embryo exhibits small local concavities or if the embryo incurs indentations through fixation and mounting. The `insideOutside` method softens the assumption of convexity by classifying points using a two-dimensional parameter space instead of requiring strict membership of a convex shape.

It has become increasingly important for developmental biologists to perform rigorous quantification of their data. Thus it is necessary to develop easy-to-deploy software that does not require high levels of programming expertise. Furthermore, benchmarking of the three mentioned classification methods has yet to be performed. Therefore, here we present the accessible `insideOutside` algorithm for the classification of interior and exterior points of embryo-like shapes. We detail the mathematics underpinning the two-dimensional parameter space used for unsupervised classification, along with accuracy testing. We then benchmark current classification methods using simulated and empirical data from pre-implantation mouse blastocysts, showing that the Convex Hull and `insideOutside` methods outperform the Ellipsoidal methods. We conclude by demonstrating that the `insideOutside` method outperforms the Convex Hull method when challenged with local surface concavities, as can be found in empirical data sets.

## Results

### The minimum distance and variance in distances from a point to the surface of a convex shape are inversely related

Here we will establish the minimum distance, and variance in distances, to the surface of a convex shape as parameters underlying the `insideOutside` algorithm. An intuitive understanding for this choice of parameters follows by considering their relationship for points at the centre, and on the surface, of a sphere.

We first consider these parameters for a point at the centre of a sphere (Fig. 2A). The minimum distance to the sphere is exactly the radius of the sphere. In fact, the distance from the centre to all other points of the sphere is identically the radius, meaning that the variance in distance to the surface of the sphere is exactly zero. Thus, minimum distance to the surface is maximized and the variance in distances is minimized for the point at the centre of a sphere. On the other hand, consider an arbitrary point on the surface of that same sphere (Fig. 2B).

**Figure 2:**
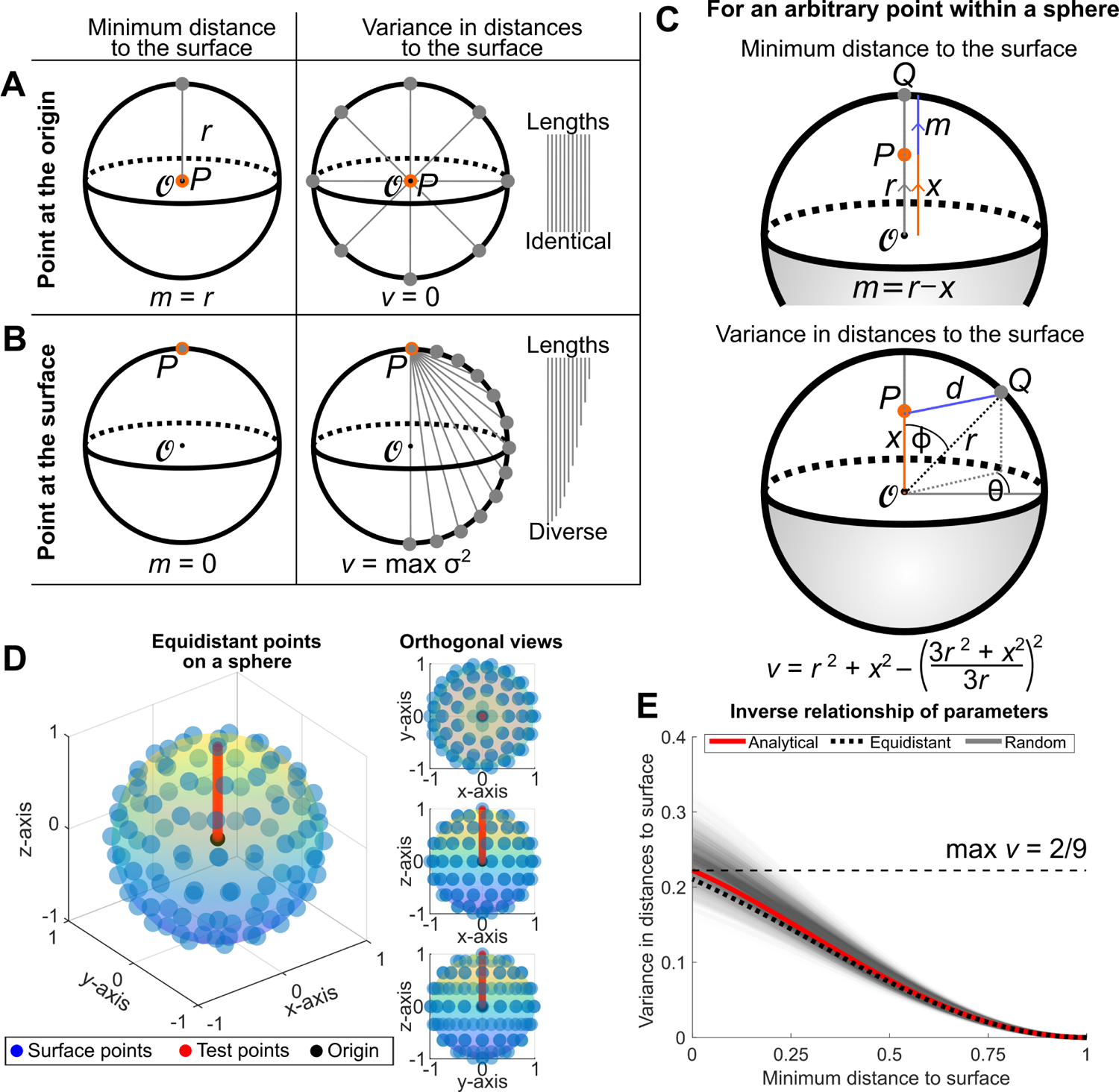
**A.** For a point *P* (orange dot) at the centre of a sphere (*O*, black dot) of radius *r*, the minimum distance from *P* to the sphere is *r* and the variance in distances is zero. **B.** For a point *P* on the surface of the sphere, the minimum distance is zero, and the variance in distances is greatest. **C.** Analytic expressions for the minimum distance, *m*, and variance in distances, *v*, from a point *P* located on/inside a sphere of radius *r* centred at the origin to the sphere. **D.** This relationship is tested for spheres that are discretized using equidistant points. 100 equidistant points (blue dots) are plotted on the unit sphere (rainbow surface). *m* and *v* are calculated for 50 test points (red dots) along the vector from the origin (black dot) to the surface point *P* = (0, 0, 1). Shown are the three-quarters view (left) and the three orthogonal views (right). **E.** The inverse relationship between *m* and *v* are shown for the continuous case of the unit sphere (see (2)) (red line) and discrete cases of 100 equidistant points on the unit sphere (Fig. 2D, black dotted line) and 100 uniform random points on the unit sphere (Fig. S1, translucent black lines, 1000 realizations).

For that surface point, the minimum distance to the surface of the sphere is exactly zero. If we then draw line segments from that point to all other points on the surface of the sphere, we see that we are drawing line segments of every length between zero and the diameter of the sphere, meaning that the original point on the surface of the sphere achieves the most diversity of line segment lengths possible for the sphere. In other words, as we will see below, a point on the surface of the sphere has the maximum variance in distances to the surface. Thus, minimum distance to the surface is minimized and the variance in distances to the surface has been maximized for any point on the surface of a sphere. We therefore arrive at an inverse relationship between the minimum distance and the variance in distances to the surface of a sphere as we move from the centre of the sphere to the surface of the sphere.

We now formalize this relationship. First, we derive the expression for the minimum distance from any point on/inside the sphere of radius *r*, centred at the origin, to the sphere. Intuitively, a point *P* on/inside the sphere, its closest point on the sphere, and the origin all lie on a straight line (Fig. 2C, top). Hence, if *P* is located a distance *x ∈* [0*, r*] from the origin, then since the distance from any point on the sphere to the origin is *r*, the minimum distance from *P* to the sphere is given by *m* = *r − x*.

Next, we derive the expression for the variance in distances from a given point on/inside the sphere to the sphere. Let *P* be located at distance *x ∈* [0*, r*] from the origin, and without loss of generality let *P* lie on the *z*-axis. Consider a point *Q* on the sphere, whose angle to the *z*-axis is given by *ϕ* (Fig. 2C, bottom). Since the distance from *Q* to the origin is *r*, by the law of cosines the distance from *P* to *Q* is given by 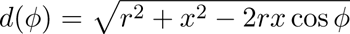. Computing surface integrals, the variance in distances from *P* to the sphere is thus given by

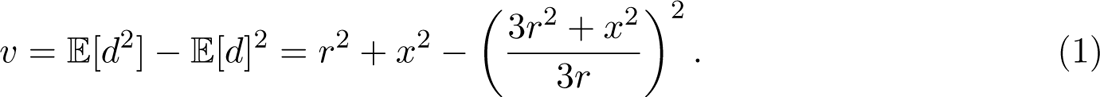

Substituting *x* = *r − m* into equation (1), and differentiating with respect to *m*, we obtain

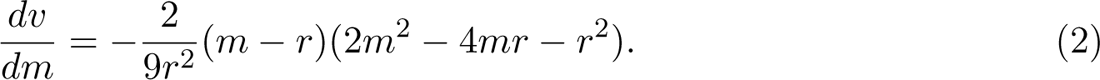

Since *m− r ≤* 0 and 2*m*^2^ *−* 4*mr − r*^2^ *<* 0 for *m ∈* [0*, r*], we have *dv/dm ≤* 0 for *m ∈* [0*, r*], hence *v* is a decreasing function of *m* for *m ∈* [0*, r*]. Thus, for points on/inside the sphere, the variance in distances is inversely related to the minimum distance to the sphere. This relationship is plotted in Fig. 2E.

The above calculations prove that the minimum distance and variance in distances to the surface are inversely related for a sphere. As an aside, we note that this relationship does not hold absolutely for arbitrary compact, convex three-dimensional shapes. Indeed, it is possible to construct examples where the centre of mass does not maximize the minimum distance to the surface, and it is this feature that hinders the inverse relationship between the minimum distance and variance in distances to the surface. Nevertheless, given a compact, convex surface which satisfies a mild roundness condition, there is a compact subset of its interior for which that the minimum distance to the boundary and the variance are inversely related. This means that the desired property holds once we are close enough to the surface, and the closer the surface resembles a sphere, the stronger the inverse relationship between the minimum distance and variance in distances to it. While a more detailed mathematical study lies outside the scope of the present study, we note that this relationship does not hold absolutely for arbitrary compact, convex three-dimensional shapes.

We conclude this section by providing numerical simulations indicating that this property remains true for piecewise linear approximations of the sphere. For this we simulate 100 points on the surface of a sphere that are spaced either equidistant (Fig. 2D) or uniformly at random (1000 realizations) (Fig. S1) (Deserno, 2004). For each discrete surface, the minimum distance and variance in distances to the surface points are calculated for 50 test points equally spaced between the centre of the sphere and a surface point. Both sets of simulations closely match the analytical solution (Fig. 2E). While this has been demonstrated for the case of the sphere and discretized derivatives of the sphere, it provides a theoretical foundation for the use of *m* and *v* in the classification of convex point-clouds. Importantly, the demonstration using discretized surfaces indicates that this relation is directly applicable to empirical data which consist of discrete points, e.g. the centroids of nuclei within an embryo.

### insideOutside: a two-dimensional decision space for classifying interior and exterior positions

Motivated by the theoretical result of the previous section, we proceed to describe an algorithm for the classification of interior and exterior points of a 3D point-cloud. The `insideOutside` algorithm (Algorithm 1) takes in a set of 3D Cartesian coordinates, *S ∈* R*^n×^*^3^ (Fig. 3Ai), and returns indexing vector *I ∈* B*^n^* with 0 indexing the inside points and 1 indexing the outside points. For pre-implantation embryos, the input data can be generated through manual nuclear segmentation in Fiji (Schindelin et al., 2012) or MATLAB’s volumeSegmenter App (Copyright 2020 The MathWorks, Inc) or through automated 3D nuclear segmentation pipelines like MINS (Lou et al., 2014), Nessys (Blin et al., 2019), or StarDist (Weigert et al., 2020). The algorithm begins by computing the Delaunay triangulation, *D*, over *S* (Fig. 3Aii). From *D*, we generate a convex hull *H* (Barber et al., 1996) (Fig. 3Aiii). Now, using *H* we can calculate the distance function, *d*(*P, H*)*, ∀P ∈ S* (Fig. 3Aiv). We calculate the minimum and variance in *d*(*P, H*)*, ∀P ∈ S*, and then scale each parameter such that the maximum is 1 and the minimum is 0 (Fig. 3Av). Note that we choose to calculate distances to all surfaces of the convex hull, not just all points generating it, to maximise accuracy and robustness of our method. Finally, hierarchical clustering by ward linkage is performed on the parameters to classify the points into two groups (Fig. 3Avi,vii). The accuracy of the `insideOutside` method classification was performed on test shapes designed to resemble the late mouse blastocyst, whose cell number ranges between 100-150 (Plusa et al., 2008), where approximately 60-70% of the cells belong to the TE (outside cells) (Fleming, 1987; Saiz et al., 2016, 2020; Morgani et al., 2018). Therefore we constructed shapes with 100 uniform random points on the unit sphere (outside) and 50 uniform random points within balls of radii between 0.01 and 1 (inside), both centred at the origin (Fig. 3B and Fig. S2). 1000 embryos were simulated and classified for each of 100 inner ball radii.

**Figure 3:**
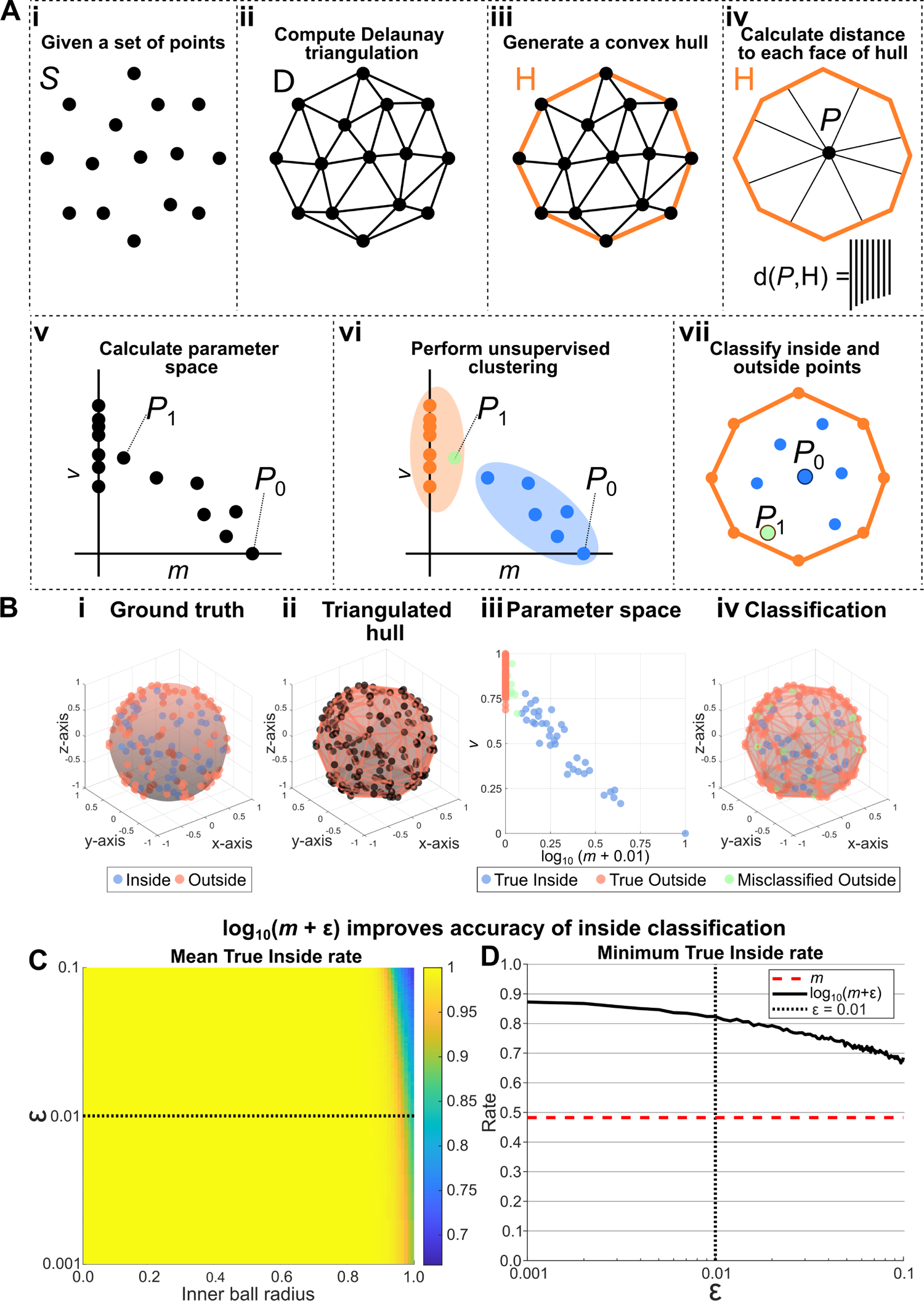
**A.** Outline of the `insideOutside` algorithm. **B.** Accuracy testing was performed on a shape constructed of 100 outside points, uniform random points on the unit sphere, and 50 inside points, uniform random points in a ball centred at *O* of radii ranging from 0.01 to 1 (see Fig. S2). **Bi-iv.** The steps of the algorithm performed on an example shape with inner ball radius of 1. **Bi.** Ground truth of inside points, blue dots enclosed by blue surface, and outside points, orange points on orange surface. **Bii.** The triangulated hull generated from making a convex hull over the Delaunay triangulation. **Biii.** The classification of points using hierarchical clustering over the calculated parameter space. Shown are True Inside points (blue), True Outside points (orange), and Misclassified Outside points (green). **Biv.** The classification mapped onto the original shape. **C-D.** Accuracy testing was performed by classifying the points (50 inside, 100 outside) of 500 shapes for each of 100 different inner ball radii for values of *ε ∈* [0.001, 0.1]. **C.** Mean True Inside rate of 500 realizations for each pair of inner ball radius and *ε*. **D.** The minimum achieved True Inside rate for each value of *ε* from the parameter sweep in **C**. The red dashed line shows the True Inside rate using the raw minimum distance and black dotted line shows the selected value of *ε*.

**Algorithm 1:**
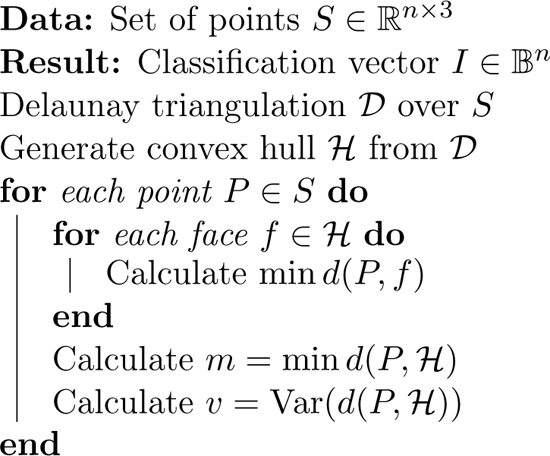
`insideOutside` takes in an *n ×* 3 matrix of Cartesian points and returns a bit vector that classifies each point as either inside, 0, or outside, 1.

### Perform unsupervised classification for two groups

Initial tests revealed near-perfect classification rates for True Outside points at all inner ball radii (Fig. S2E). There was, however, a significant drop in the True Inside classification rate at an inner ball radius of 0.82 where the True Inside classification rate dropped below 0.99 with minimum rate of 0.48 ± 0.17 (mean ± standard deviation) at an inner ball radius of 1. To improve the True Inside classification rate, we modified the parameter space by taking the log of *m*, which results in a greater separation of inside and outside points along the minimum distance axis (Fig. S2F). Simulations bear out marked improvements in True Inside classification rates with no detriment to True Outside classification rates. The resulting True Inside classification rates do not drop below 0.99 until an inner ball radius of 0.94 and achieve a minimum of only 0.82 ±0.09 at an inner ball radius of 1.

We extended this analysis to explore the effects on True Inside classification rate by adding different values of a small parameter, *ε*, to the minimum distance to the surface (Fig. 3C). We observe very high True Inside classification rates (>0.95) for the vast majority of inner ball radii. For all values of *ε* assessed, a decrease in True Inside classification rate is only observed for inner ball radii near one. Indeed, we see that the worst case True Inside classification rates for each *ε* are obtained for inner ball radii at, or very near, one (Fig. 3D). While these worst case True Inside classification rates range between 0.67 and 0.87, each *ε* outperforms the raw minimum distance of 0.48. We therefore conclude that so long as *ε* is sufficiently small, the exact value has little impact on the classification rates. Thus we have established our algorithm using the parameters of [log_10_(*m* + *ε*)*, v*], for *ε* = 0.01, and we now proceed to challenge this method with empirical data.

### insideOutside and Convex Hull methods outperform the Ellipsoidal methods when classifying cells of the mouse blastocyst

In this section we set out to show that the `insideOutside` method can successfully classify the nuclei of real-world samples by using previously quantified mid-blastocysts mouse embryos (Stirparo et al., 2021) (Fig. 4A). We initially assess different unsupervised clustering mehtods which can be used in `insideOutside` and how well the minimum distance and variance of distances to the surface relationship is able to classify empirical data. Next we assess how well the individual parameters, minimum distance and the variance of distances to the surface, are able to classify these data. We finally benchmark our method against the three other methods (Naïve Ellipsoidal, RANSAC Ellipsoidal (Lou et al., 2014), and Convex Hull (Forsyth et al., 2021)) with these same embryos. SOX2 staining, which can be used to mark all nuclei of the early ICM (Wicklow et al., 2014), was used as the ground truth, where SOX2 positive nuclei indicate inside nuclei and SOX2 negative nuclei indicate outside nuclei. SOX2 positive/negative status was determined through statistical inference (Gaussian mixture modelling) with 758 cells from 14 embryos (Fig. 4B,C). We then used the SOX2 ground truth (Fig. 4D) to calculate the True Inside and True Outside rates for the four classification methods(Fig. 4E).

**Figure 4:**
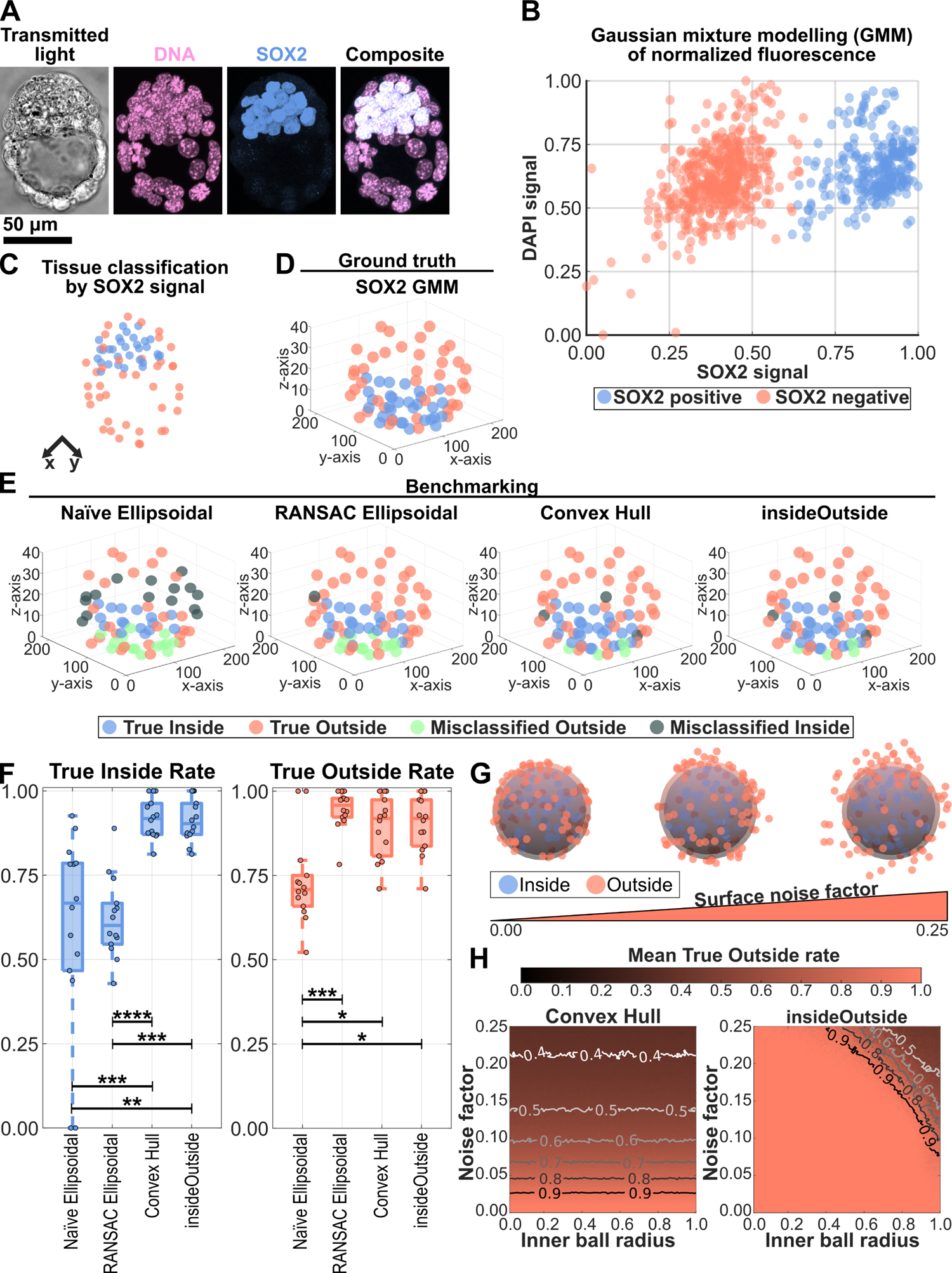
**A.** Confocal images of a mid-blastocyst stained for DNA and early ICM marker SOX2. A single slice is shown for transmitted light and maximum intensity projections are shown for fluorescence images. **B.** Gaussian mixture modelling (GMM) was performed on the SOX2 nuclear signal of 758 cells from 14 embryos to classify SOX2 positive (blue) and negative (orange) nuclei. Nuclear signal was normalized by nuclear volume, log_10_ transformation, and re-scaling to the interval [0,1]. **C.** GMM classification of cells applied to the embryo from **A**. **D.** SOX2 GMM classification of embryo from **A** shown in three-quarters view. SOX2 GMM classification was used as the ground truth for methods benchmarking. Shown are True Inside (blue) and True Outside (orange) cell classification. **E.** Classification of embryo from **A** by the Naïve Ellipsoidal, RANSAC Ellipsoidal (Lou et al., 2014), Convex Hull (Forsyth et al., 2021), and `insideOutside` (Stirparo et al., 2021) methods shown in three-quarter view. Shown are True Inside (blue), True Outside (orange), Misclassified Outside (green; inside classified as outside), and Misclassified Inside (gray; outside classified as inside) nuclei. **F.** Classification rates for True Inside (left, blue) and True Outside (right, orange). Kruskal-Wallis Test, p-values: *, 0.05 *≥* p > 0.01; **, 0.01 *≥* p > 0.001; ***, 0.001 *≥* p > 0.0001; ****, 0.0001 *≥* p. All other pairwise relationships were not significant with p-values *≥* 0.63. **G.** Increasing levels of noise were added to the surface points of the test shape to simulate increasing local surface concavities. **H.** The mean True Outside rate (orange scale) is shown over the parameter space of inner ball radius (100 radii between 0.01 and 1) versus noise factor (100 levels between 0 and 0.25) for the Convex Hull (left) and `insideOutside` (right) methods. 100 test shapes were classified for each parameter pair. Additional contour lines are shown to delineate drops in classification rate.

First, we assessed different unsupervised clustering methods on classification rates (Fig. S3A). The methods we assessed were K-means clustering, hierarchical clustering, spectral clustering, and DBSCAN. No difference was observed in either the True Inside classification rate (means ranging between 0.91 and 0.93, p-values > 0.05, Kruskal-Wallis) or the True Outside classification rate (p-values > 0.05, Kruskal-Wallis). However, the spectral clustering method exhibited a larger variance than the other three methods and a lower mean rate of 0.79, while the mean rates for the other three methods ranged between 0.89 and 0.90. We therefore propose that any of K-means clustering, hierarchical clustering, or dbscan are sufficient for unsupervised clustering. However, because DBSCAN requires additional user defined parameters, we would only suggest K-means clustering or hierarchical clustering. Finally, we select hierarchical clustering for insideOutside as the preferred method as it tends to better classify highly eccentric distributions, such as the ones generated from these data (Fig. S2C).

Next, we see from simulations that there is a roughly linear relationship between the minimum distance to the surface, *m*, and variance of distances to the surface, *v*, for points away from the surface and centre of a sphere (Fig. 2E). Therefore, we wanted to determine if one of the parameters was sufficient for classification of empirical data, or if it is the case that both parameters are needed to improve classification in the nonlinear regions that are towards the surface and centre of embryos. To that end, we examined the ability to classify nuclei using only a univariate parameter space of either log_10_(*m* + 0.01) or *v* against the bivariate parameter space of [log_10_(*m* + 0.01)*, v*] (Fig. S3B. We see that *v* alone is a relatively poor classifier for both inside nuclei (rate = 0.69 ± 0.21, mean ± standard deviation; N = 14 embryos) and outside nuclei (rate = 0.65 ± 0.15). Whereas log_10_(*m* + 0.01) and [log_10_(*m* + 0.01)*, v*] correctly classified nuclei with identical rates (rate = 0.91 ± 0.06) above that of *v* alone (p-value of 0.0027, Kruskal-Wallis). Again log_10_(*m* + 0.01) (rate = 0.895 ± 0.092; p-value of 0.004) and [log_10_(*m*+0.01)*, v*] (rate = 0.904 ± 0.086; p-value of 0.002) yielded better rates for outside nuclei classification rates compared with *v*. Although, outside classification rates for log_10_(*m* + 0.01) and [log_10_(*m* + 0.01)*, v*] were not significantly difference (p-value of 0.98), [log_10_(*m* + 0.01)*, v*] outperformed log_10_(*m* + 0.01) for 3 out of 14 embryos. This leads us to conclude that the bivariate parameter space does indeed provide an advantage in classifying outside nuclei in some instances.

With `insideOutside` in hand, we now benchmark our method against the three other methods. For inside nuclei classification (Fig. 4F,left), we find that both the Convex Hull (rate = 0.92 ± 0.06, mean ± standard deviation; N = 14 embryos) and `insideOutside` (rate = 0.91 ± 0.06) methods outperform the Naïve Ellipsoidal method (rate = 0.59 ± 0.29; p-values of 6.8 *×* 10*^−^*^4^ and 1.0 *×* 10*^−^*^3^, respectively, Kruskal-Wallis) and RANSAC Ellipsoidal method (rate = 0.62 ± 0.12; p-values of 8.1 *×* 10*^−^*^5^ and 1.3 *×* 10*^−^*^4^, respectively). The Naïve and RANSAC Ellipsoidal methods show no difference in ability to classify inside nuclei (p-value = 0.96). Similarly, the Convex Hull and `insideOutside` methods show no difference in ability to classify inside nuclei (p-value = 0.99, Kruskal-Wallis). These rates are summarized in the table in Fig. S3C.

For outside nuclei classification (Fig. 4F,right), we see that the Naïve Ellipsoidal method (rate = 0.73 ± 0.13) underperforms compared to the RANSAC Ellipsoidal method (rate = 0.95 ± 0.06, p-value = 4.9 *×* 10 *^−^*^4^), Convex Hull method (rate = 0.89 ± 0.09, p-value = 0.031), and `insideOutside` method (rate = 0.90 ± 0.08, p-value = 0.019). However, no difference is seen between the RANSAC Ellipsoidal, Convex Hull, and `insideOutside` methods (all pair-wise p-values *≥* 0.63). Thus, the Naïve Ellipsoidal method shows the lowest rates of classification for the both inside and outside points. While the RANSAC Ellipsoidal method only underperforms in classifying interior points. These data together show that ellipsoid fitting has the least satisfactory classification accuracy.

### insideOutside outperforms the Convex Hull method in classification of outside points

While both the `insideOutside` and Convex Hull methods perform comparably on the empirical blastocysts, the Convex Hull method holds a systematic error of misclassifying outside points as inside points when there are minor concavities at the surface. We highlight this feature in two head-to-head comparisons of the `insideOutside` and Convex Hull methods. First we simulate segmentation/imaging errors by leave-K-out sub-sampling of the empirical embryos. Then we conclude with the introduction of small local surface concavities to simulated test shapes.

We begin by simulating segmentation/imaging errors by k-fold cross validation sub-sampling of the empirical embryos for k values between 1 and 15. This reveals that the Convex Hull method outperforms `insideOutside` in classifying inside points in 16.44% of of sub-sampled embryos (Fig. S3D). When considering outside classification, `insideOutside` performs better in 12.98% of sub-sampled embryos while the Convex Hull method performs better in 0.06% of sub-sampled embryos (Fig. S3E). We then asked how sensitive these methods are to perturbations relative to the non-subsampled embryos. We found a decrease in inside classification rates relative to the non-subsampled classfication for both Convex Hull method (39.86% of samples) and `insideOutside` (40.71% of samples) (Fig. S3F). Conversely, the an increase in outside classification rates relative to the non-subsampled embryos was exhibited for both Convex Hull method (31.37% of samples) and `insideOutside` (27.70% of samples) (Fig. S3G).

We now show the systematic nature of this misclassification by emulating increasing levels of local surface concavities via the introduction of increasing levels of normally distributed random noise to the surface points of the test shapes (Fig. 4G). We then compute the classification rates over the parameter space of inner ball radius (100 radii between 0.01 and 1) and noise factor (100 levels between 0 and 0.25) for 100 shapes (Fig. 4H and Fig. S4A). The Convex Hull method shows a uniform decrease of True Outside classification rates across all inner ball radii for increasing levels of noise, eventually dropping below a rate of 0.4 around a noise factor greater than 0.2. The `insideOutside` method does not display this uniform decrease of True Outside classification rates, instead maintaining a rate of greater than 0.9 for the majority of the parameter sets tested. The `insideOutside` method only begins to lose accuracy when both the inner ball radius and noise factor become large. Surface concavities have negligible effects on the classification rates of inside points for both methods (Fig. S4B,C).

## Discussion

Motivated by the need to accurately classify cells of mouse embryos based on their spatial position alone, we present `insideOutside`, an accessible algorithm for the classification of interior and exterior points of a three-dimensional point-cloud. We established an inverse relationship between the minimum distance and variance in distances from a point to a ‘typical’ convex shape’s surface. We then harnessed this inverse relationship to build an algorithm which allows for faithful classification of interior and exterior points by hierarchical clustering. We then proceeded to benchmark our method against three other methods, Naïve Ellipsoidal, RANSAC Ellipsoidal (Lou et al., 2014) and Convex Hull (Forsyth et al., 2021), finding that the `insideOutside` method was as reliable, or better, at classifying nuclei of the pre-implantation mouse embryo. We demonstrated that the `insideOutside` method has greater accuracy than the Convex Hull method in classifying exterior points when challenged with both segmentation/imaging errors and local surface concavities. Finally, we have packaged the algorithm as freely available stand-alone MATLAB and Python implementations.

We have shown that the Convex Hull and `insideOutside` methods both outperform the Naïve and RANSAC Ellipsoidal methods in the classification of interior nuclei of pre-implantation mouse embryos, while the Naïve Ellipsoidal method underperforms the other three methods when classifying exterior nuclei. In all four methods we find that instances of misclassification are highest where the ICM is in contact with the TE (Fig. 4E). This shows that ellipsoid fitting methods (Naïve and RANSAC) are not as suitable in classifying interior and exterior points as convex hull based methods (Convex Hull and `insideOutside`).

Moreover, we have shown through simulation that the `insideOutside` method is more accurate than the other methods when challenged with surface concavities. This is particularly important for classifying model systems whose exterior points exhibit high levels of noise. Such noise can be biological in nature, e.g. due to variability in nuclear height along the apicobasal axis of columnar epithelia; or technical, e.g. due to segmentation errors.

These findings speak to the appropriateness of each method. There may be instances when the user has a large number of points in a low noise situation where exterior points should be strictly classified as belonging to the surface. In such cases the Convex Hull method is most appropriate. Alternatively, the user may want to soften this condition in the case of a small number of points in a high noise situation, e.g. the pre-implantation mouse embryo. Here, the `insideOutside` method would be most useful, as it performs the best in such high noise situations. While the Ellipsoidal method has proved useful in identifying unique embryos from images with many embryos (Lou et al., 2014), we would not recommend the Ellipsoidal method for classifying the nuclei of those embryos. Finally, we note that all of the classification methods considered in the present study take on the order of 10*^−^*^4^ to 10*^−^*^3^ s to implement for a typical mouse embryo. As such, relative computational cost is unlikely to be a factor unless analysing unrealistically large numbers of embryos.

There is scope for further refinement of the `insideOutside` algorithm. This could come by way of incorporating more information about the segmented nuclei, e.g. making use of nuclear aspect ratio and not just nuclear centroid. Additional parameters could also be introduced to the parameter decision space. IVEN has made use of number-of-neighbours, calculated from the Delaunay triangulation, in downstream spatial analysis. The addition of number-of-neighbours to the classification space may aid in better discrimination of interior and exterior points, especially in the problem case where the ICM meets the TE. More generally, extensions to our algorithm could leverage alternative approaches to improve robustness of classification, such as those based on deep learning (Ounkomol et al., 2018; Christiansen et al., 2018).

We have sought to make this method, and future methods, easy-to-deploy for biological and life scientists, as there is increasing need for them to perform rigorous quantification of their data. Development of the `insideOutside` method in Stirparo et al. (2021) was born of a need to refine the original MINS classification method and was driven by collaboration between experimentalists and theoreticians. While there is no expectation for experimentalists to do methods development, there is expectation that they should be able to use these methods, thus empowering future work. Both MINS and IVEN share this ethos of empowering experimentalists in the journey of data analysis. However, the `insideOutside` method is provided as a stand-alone script, whereas the classification methods in MINS and IVEN are members of a larger software package. This means that the `insideOutside` method has greater flexibility in use and migration to other programming languages. Critically, the stand-alone nature of the `insideOutside` method lends itself to incorporation into other software pipelines. For example, the `insideOutside` method could be incorporated as an additional classification method into either MINS or IVEN, as both packages have MATLAB implementations.

Finally, other use cases for the `insideOutside` method include other mammalian organisms that undergo the process of blastocyst formation (humans, non-human primates, other rodents, ungulates, etc.). It also has use for certain organoid systems, such as quantifying the level of cell sorting in ICM organoids (Mathew et al., 2019). And while the `insideOutside` method was motivated by the need to discriminate between the ICM and the TE in the pre-implantation blastocyst, it remains a general method for classifying the interior and exterior points of a point-cloud. This means it has extensibility to any data of this description. This includes the organization of transcription factor clusters from single-molecule localization microscopy (Liu et al., 2014), the pattern of RNA transcripts acquired through seq-FISH (Lohoff et al., 2022), and the relationship of genomic loci within the nucleus as determined by single-cell Hi-C structures (Stevens et al., 2017).

## Materials and Methods

### Embryo collection and bright-field imaging

Embryos were obtained from natural mating, detection of a copulation plug in the morning was used as confirmation of successful mating and indicated embryonic day (E) 0.5. 8-cell and compacted morula embryos were flushed from the oviduct at E2.5 and E3.0, respectively, and mid and late blastocysts were flushed from the uterine horns at E3.5 and E4.5, respectively, using M2 medium (Sigma-Aldrich, M7167). This research has been regulated under the Animals (Scientific Procedures) Act 1986 Amendment Regulations 2012 following ethical review by the University of Cambridge Animal Welfare and Ethical Review Body. Use of animals in this project was approved by the ethical review committee for the University of Cambridge, and relevant Home Office licences (Project licence No. 80/2597 and No. P76777883) are in place. Bright-field images were taken on a Leica DMI4000B microscope.

### Quantitative immunofluorescence of embryos

Quantitative immunofluorescence data was originally published in Stirparo et al. (2021). In brief, embryos were fixed in paraformaldehyde, stained for DNA and SOX2, and imaged using confocal microscopy. Embryo nuclei were segmented and each nucleus’s total fluorescence (sum of pixel values), volume, and centroid were quantified using MINS (Lou et al., 2014).

### Accuracy testing designations and rates

For accuracy testing, each point was assigned one of four designations: true inside, misclassified inside, true outside, or misclassified outside. A designation of “true inside” means the ground truth of the point and the classification of the point were both “inside”. A designation of “misclassified inside” means the ground truth of the point was “inside” and the classification of the point was “outside”. The same logic applies to the “outside” designations.

Two rates were calculated in the accuracy testing: true inside rate and true outside rate. The “true inside rate” was calculated as the number “true inside” points divided by the number of “total inside” points determined by the ground truth,

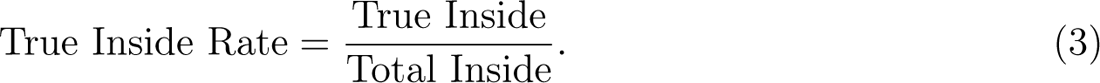

The same logic applies to the “true outside rate”. Note that our terminology is based on biological intuition and represents a departure from standard machine learning nomenclature: “true inside rate” corresponds to the sensitivity of the inside classification, and similarly for “true outside rate”.

### Simulating segmentation error

Segmentation error was simulated by *k*-fold cross validation sub-sampling, *k ∈* [1, 15], of the 14 different embryos. For one-fold cross validation sub-sampling, each nucleus was left out once to build the sample (E.g. For an embryo with 35 nuclei, there would be 35 samples drawn). For *k*-fold cross validation sub-sampling with *k >* 1, we randomly chose 100 samples with replacement from each of the 14 embryos, for a total of 1400 sub-sampled embryos for each value of *k*.

### Code availability

The `insideOutside` algorithm and all code used in this manuscript to perform simulations, analysis, and benchmarking are written in MATLAB (2021a) and are freely available at https://github.com/stanleystrawbridge/insideOutside under the GNU General Public License v3.0. A Python implementation of the `insideOutside` algorithm is provided in the same repository.

## Acknowledgements

The authors thank the members of the Fletcher and Nichols groups for their helpful feedback in the preparation of the manuscript, especially Ian Groves and Lawrence Bates. Collaboration between the Fletcher and Nichols groups was made possible through a Company of Biologists Travelling Fellowship awarded to SES (DEVTF-180513). SES also acknowledges a Sir Henry Wellcome Postdoctoral Fellowship (224070/Z/21/Z). AGF acknowledges support from the Biotechnology and Biological Sciences Research Council (BB/V018647/1 and BB/R016925/1).

## Author contributions

SES conceived and developed the `insideOutside` algorithm. ECS collected bright field images of pre-implantation mouse embryos. SES, ANF and AGF performed mathematical analyses. AK performed embryo immunostainings and segmentation used for methods benchmarking. SES and AGF performed simulations and computational analyses. SES, AGF, and JN wrote the manuscript.

## Supplemental material

**Figure S1:**
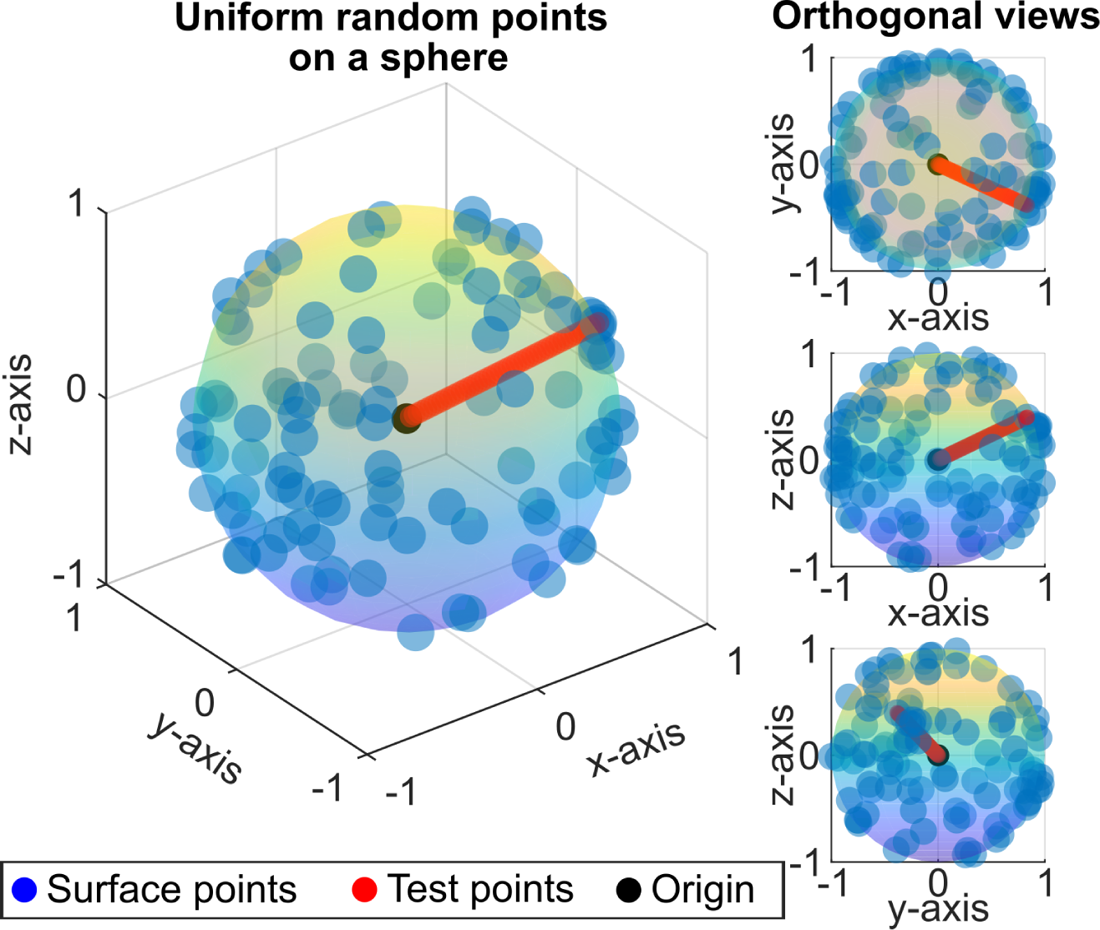
Example of 100 random points (blue dots) drawn uniformly from the unit sphere (rainbow surface). The minimum distance to the surface, *m*, and variance in distances to the surface, *v*, are calculated for 50 test points (red dots) along the vector from the origin (black dot) to a random point on the surface. Shown are the three-quarters view (left) and three orthogonal views (right).

**Figure S2:**
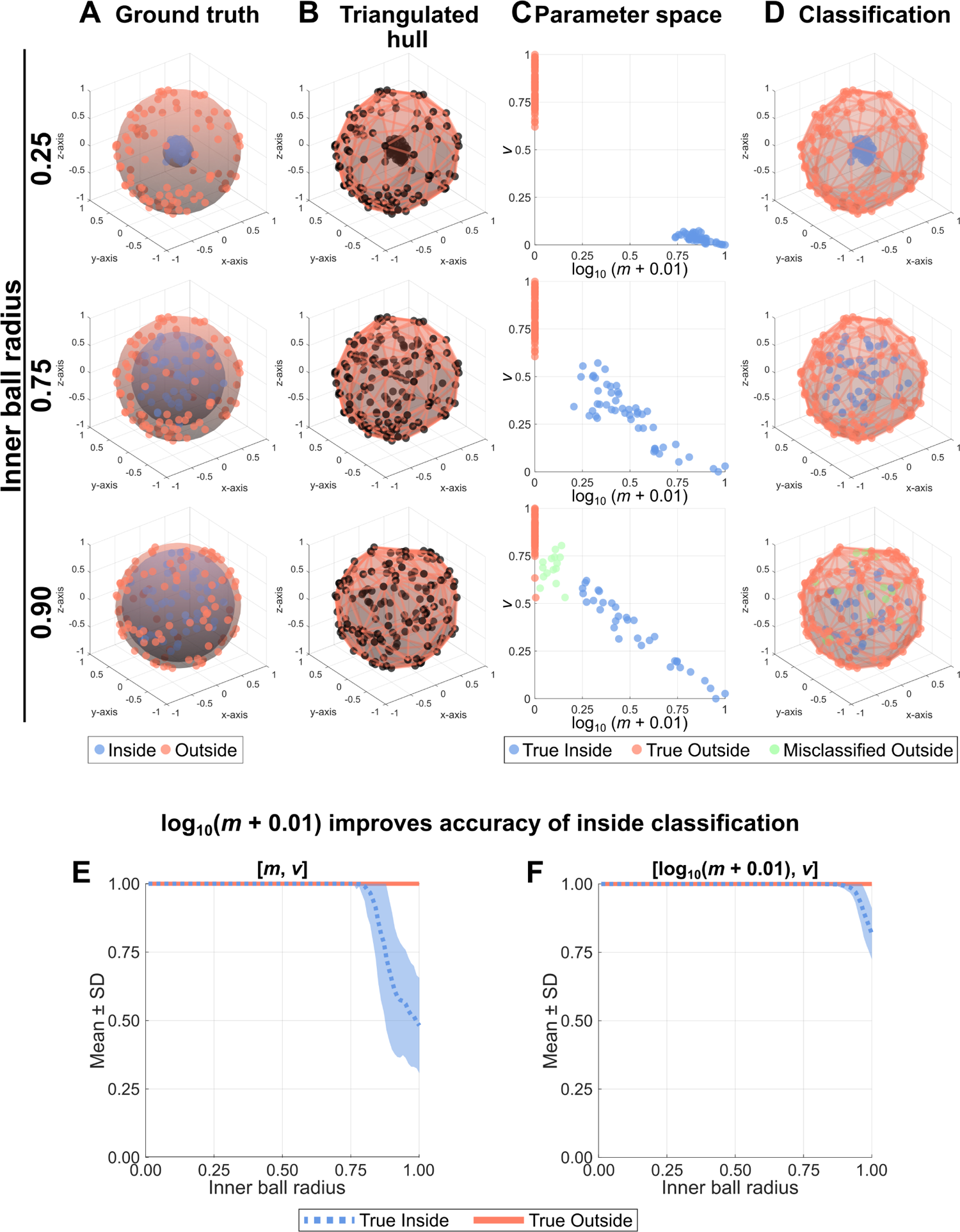
Example shapes (50 inside points, 100 outside points) with different inner ball radii used for accuracy testing. **A.** Ground truth of inside points (blue dots enclosed by blue surface) and outside points (orange points on orange surface). **B.** The convex hull generated by the Delaunay triangulation. **C.** Classification of points using hierarchical clustering over the calculated parameter space. Shown are True Inside points (blue), True Outside points (orange), and Misclassified Outside points (green). **D.** The classification mapped onto the original shape. **E-F.** Accuracy testing was performed by classifying the points of 1000 shapes for 100 different inner ball radii. The mean True Inside rate (blue dotted line) is shown with standard deviation (blue filled region) and the mean True Outside rate (orange solid line) is shown with standard deviation (orange filled region). **E.** Accuracy test for the parameters *m* and *v*. **F.** Accuracy test for the parameters log_10_ (*m* + 0.01) and *v*.

**Figure S3:**
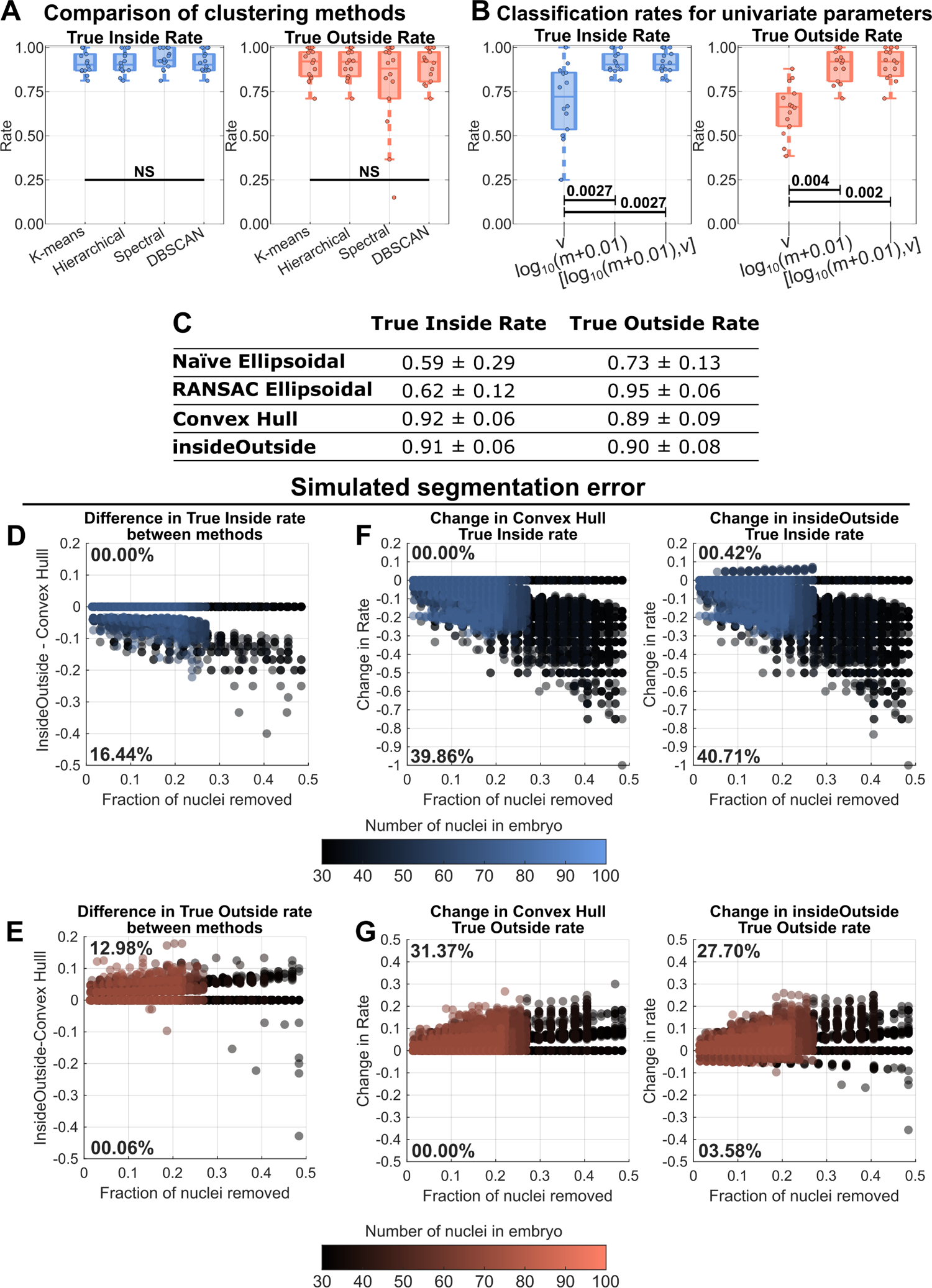
**A.** Classification comparison for different unsupervised clustering methods (NS = not significant, p-values > 0.05, Kruskal-Wallis Test). **B.** Classification comparison using a univariate parameter space of mean distance to surface, log_10_(*m* + 0.01), or variance in distances to surface, *v*, or the bivariate parameter space [log_10_(*m* + 0.01)*, v*] (p-values, Kruskal-Wallis Test). **C.** Summary of classification rates for different methods. Shown are the mean and standard deviation. **D-G.** Assessing the effects of simulated segmentation error on classification rates. **D-E.** The difference in classification rates between the indieOutside and the Convex Hull Methods for **(D)** inside nuclei and **(E)** outside nuclei. **F-G.** The change in classification rate from the non-subset embryos for indieOutside and the Convex Hull Methods for **(F)** inside nuclei and **(G)** outside nuclei.

**Figure S4:**
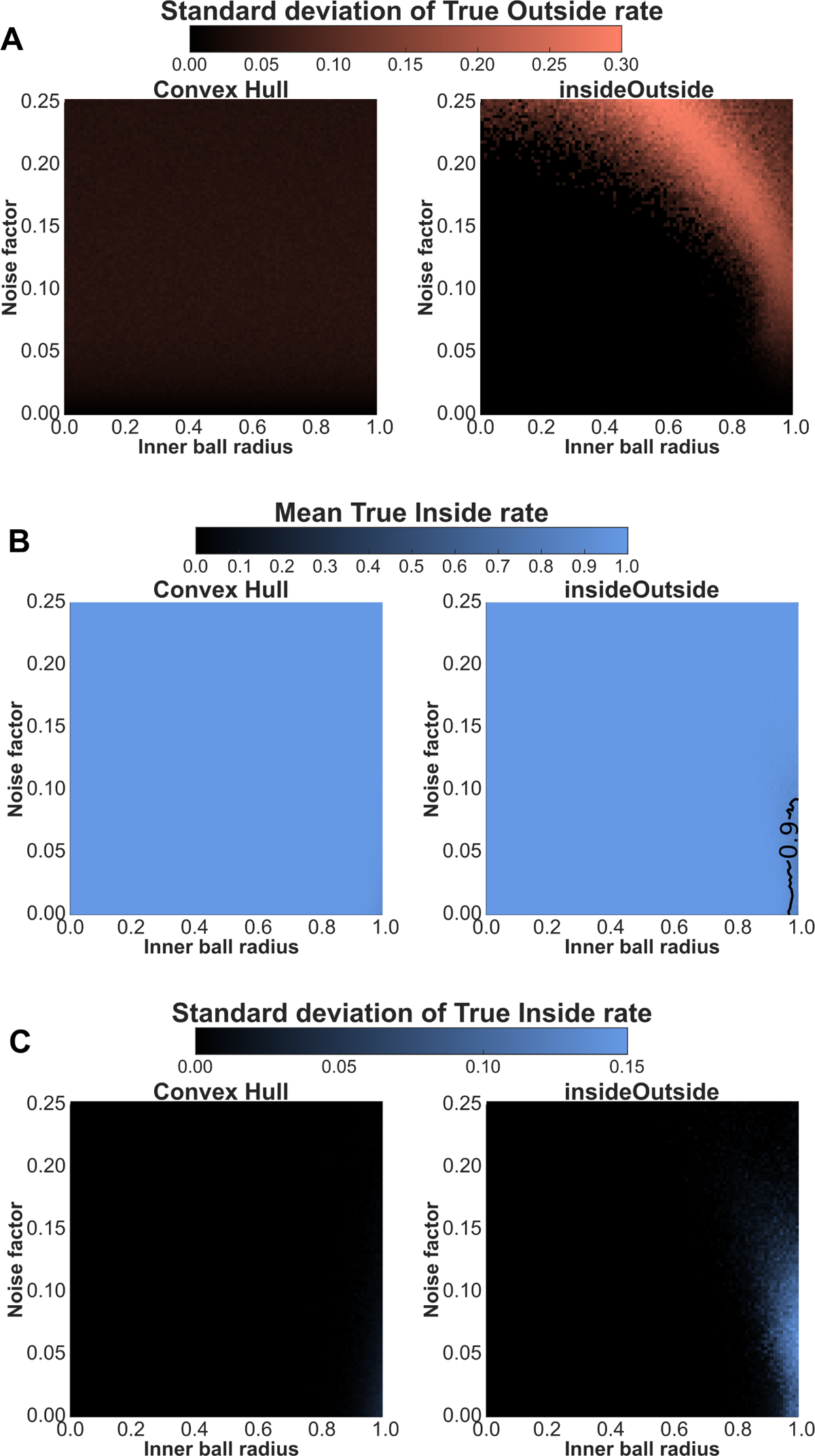
Classification rates shown over the parameter space of inner ball radius (100 radii between 0.01 and 1) versus noise factor (100 levels between 0 and 0.25) for the Convex Hull (left) and `insideOutside` (right) methods. **A.** The standard deviation of the True Outside rate. **B.** The mean True Outside rate. Additional contour lines are shown to delineate drops in classification rate. `insideOutside` rate is > 0.9 everywhere except when the inner ball radius is close to 1 and the noise factor is small. **C.** The standard deviation of the True Inside rate.

